# Seawater salt-trapped *Pseudomonas aeruginosa* survives for years and gets primed for salinity tolerance

**DOI:** 10.1101/649152

**Authors:** Hamouda Elabed, Enrique González-Tortuero, Claudia Ibacache-Quiroga, Amina Bakhrouf, Paul Johnston, Kamel Gaddour, Jesús Blázquez, Alexandro Rodríguez-Rojas

## Abstract

**Background:** In nature, microorganisms have to adapt to long-term stressful conditions often with growth limitations. However, little is known about the evolution of the adaptability of new bacteria to such environments. *Pseudomonas aeruginosa*, an opportunistic pathogen, after natural evaporation of seawater, was shown to be trapped in laboratory-grown halite crystals and to remain viable after entrapment for years. However, how this bacterium persists and survives in such hypersaline conditions is not understood.

**Results:** In this study, we aimed to understand the basis of survival, and to characterise the physiological changes required to develop salt tolerance using *P. aeruginosa* as a model. Several clones of *P. aeruginosa* were rescued after fourteen years in naturally evaporated marine salt crystals. Incubation of samples in nutrient-rich broth allowed re-growth and subsequent plating yielded observable colonies. Whole genome sequencing of the *P. aeruginosa* isolates confirmed the recovery of the original strain. The re-grown strains, however, showed a new phenotype consisting of an enhanced growth in growing salt concentration compared to the ancestor strain. The intracellular accumulation of K^+^ was elicited by high concentration of Na^+^ in the external medium to maintain the homeostasis. Whole transcriptomic analysis by microarray indicated that seventy-eight genes had differential expression between the parental strain and derivative clones. Sixty-one transcripts were up-regulated, while seventeen were down-regulated. Based on a collection of single-gene knockout mutants and gene ontology analysis, we suggest that the adaptive response in *P. aeruginosa* to hyper-salinity relies on multiple gene product interactions.

**Conclusions:** The individual gene contributions build up the observed phenotype, but do not ease the identification of salinity-related metabolic pathways. The long-term inclusion of *P. aeruginosa* in salt crystals primes the bacteria, mediating a readjustment of the bacterial physiology to growth in higher salt concentrations. Our findings provide a starting point to understand how *P. aeruginosa*, a relevant environmental and pathogenic bacterium, survives to long-term salt stress.

## Background

In their natural environments, microbes often have to cope with stressful conditions. The limitation of nutrients, intense competition for resources and a variety of abiotic stresses such as radiation, temperature, pH, oxygen-derived radicals, antibiotics and high osmolarity are commonly experienced by bacteria [1].

Reports on the extreme longevity of microbes in salt are controversial [2]. Hypersaline environments have been a significant reservoir for the long-term evolution of specially adapted microorganisms [3]. Additionally, saline environments may aid the survival of microorganisms, protecting them from desiccation by trapping the cells in fluid inclusions, a phenomenon that occurs in salt crystals upon evaporation [4]. Several studies on ancient microbes are consistent with laboratory experiments and studies on other modern surface halite deposits, which suggest that microorganisms persist inside fluid inclusions in halite for many year [5–7].

In a previous study, *Pseudomonas aeruginosa* cells were shown to get trapped in fluid inclusions pockets of saturated brine in laboratory-grown halite crystals and to remain viable after entrapment [5]. The ability of *P. aeruginosa* to colonise and thrive in myriad environments correlates with its relatively large genome and genetic complexity [8]. An exceedingly high number of assigned open reading frames are transcriptional regulators or members of two-component regulatory systems in comparison to other bacteria [9]. This large proportion of regulatory genes also facilitates *P. aeruginosa* adaptability and sensing diverse environmental stresses [8–11]. Potassium is the major intracellular cation in bacteria and plays an important role to maintain homeostasis. In osmotic conditions, bacterial cells accumulate K^+^ by a number of different transport systems that vary in kinetics, energy coupling, and regulation [12].

However, despite advances in the understanding of the immediate response to hyperosmotic shock in *P. aeruginosa* [13], the physiological mechanisms that allow the bacterial persistence in highly saline environments are still poorly understood. Molecular basis of this persistence may be of great interest to both clinical and environmental microbiology. In the present study, we assess the phenotypic and genotypic changes of *P. aeruginosa* ATCC 27853 after fourteen years of entrapment in seawater salt crystals to characterise the required physiological changes that allow salt tolerance.

## Results

### Evaluation of fitness in different salt conditions

In this work, we study the effects of long-term incubation in extremely salty conditions on *P. aeruginosa* using microarrays and salt-tolerance assays. After fourteen years of inclusion in evaporated seawater (37 g/l of salts), different clones of *P. aeruginosa* were recovered and cultivated. The revitalisation of the culture in nutrient broth at 37°C rendered bacterial suspensions that reached an average of OD_600nm_: 0.9±0.06, after 48 hours of incubation. The ancestor strain (T0 or control) and its derivative 48-hour clones (T48), did not show significant differences in growth rate when cultured with NaCl 8.5 mM (normal concentration of NaCl in DM medium (Table 1). However, the final OD was significantly higher in the recovered T48 strain. This implies a growth advantage in the stationary phase demonstrating the adaptability of the recovered cells to extremes conditions such as starvation. After the long period in salt crystals, supposing the selection of more adapted mutants, we also cultured the bacteria on increasing concentrations of NaCl. The variants T48 showed an improved growth rate at concentrations of 250 or 500 mM (Table 1). At 1M, the ancestor strain was not able to grow, whereas T48 clones reached the highest optical density of all conditions (Figure 1). These first results suggested that derivative T48 clones acquired the ability to thrive in high-salt environments, even at NaCl concentrations that were restrictive for the original strain. However, no significant differences were found when comparing growth rates (*r*) between T0 and T48 variants at other evaluated NaCl concentration (Table 1). Moreover, the addition of 100 mM KCl to bacterial cultures of the T0 strain, inhibited by 1M of NaCl, restored the growth of this strain and allowed T48 strain to growth even at 2M NaCl (Figure 2), indicating that growth inhibition not only depends on salt concentration but also on the composition of growth media. The tolerance to NaCl is then influenced by the level of KCl or maybe the ability of the cell to control K^+^ transport. Hence, our expectations were to find the role of K^+^ and Na^+^ transporters or regulators in *P. aeruginosa* to long-term hypertonic conditions. All these observations demonstrate that after incubation of *P. aeruginosa* in seawater crystals for a long period (14 years), the cells adapted and became more tolerant to higher salt concentrations.

### Whole genome sequencing after recovery from salt

To characterise possible genomic adaptations to salt in *P. aeruginosa*, we sequenced five independent clones and the reference strain using a whole genome sequencing approach based on the Illumina MiSeq platform. Seven non-synonymous substitutions were found in the sequenced clones. These mutations were present in one aromatic amino acid transporter, prepilin-type N-terminal cleavage/methylation domain-containing protein, FHA domain-containing protein, hybrid sensor histidine kinase/response regulator and two hypothetical proteins. In addition, three SNPs resulting in a synonymous substitution (c.795G>C p.Arg265Arg; c.34T>C p.Leu12Leu and c.54C>T p.Ser18Ser) respectively in DNA polymerase III subunit beta and in two hypothetical proteins were also detected by WGS (Table S1). Three other mutations were identified in intergenic sites. The products of the genes with non-synonymous mutations do not seem to contribute in salt stress response. Overall, the lack of convergence in the mutations makes difficult to assign any effect to these SNPs. Suggesting that the changes on salt resistance in the studied strain were probably associated with adaptive response based on changes in global gene expression.

### Transcriptome profiling of salt-trapped *P. aeruginosa*

As the phenotype of T48 clones cannot be easily explained by mutations, differences between T0 and T48 may be due to physiological changes that remain after the recovery from the seawater salt crystals. To investigate the differential gene expression between the wild-type T0 and its salt-tolerant derivate T48, transcriptome analysis by microarrays rendered 78 genes with significant changes in their expression level. From these 78 genes, 25 are genes related to cellular metabolism, 18 are associated to virulence factors, 14 are hypothetical proteins, 11 are associated to transporters and regulatory peptides, 4 are membrane proteins, 4 are implicated in post translational modification and 2 are chaperones and heat shock proteins (Table 2). A global view of all differentially expressed genes is presented in the MA plot in the Figure S1.

From the 25 genes encoding enzymes or proteins implicated in the cellular metabolism which are differentially expressed between T0 and T48, 20 genes are up-regulated and five are down-regulated. When the 18 genes associated with expression of virulence factors are analysed, all Type III Secretion System (T3SS) proteins and two cytotoxin secretion factor exoenzymes, ExoS (PA3841) and ExoT (PA0044) are up-regulated (Table 2). A remarkable increase of the expression of H^+^ transport T3SS ATPase (*pscN* – PA1697) was also observed in T48 strains. Interestingly, its product can be responsible for Na^+^ extrusion in *P. aeruginosa*. A substantial homeostatic capacity is necessary for adaptation and tolerance to a change in the external environment.

There are eight genes encoding transporters that were up-regulated in T48. Except for the genes related to the symport of Na^+^/Alanine/Glycine (PA2252), transport of sulphate (*cysW*– PA0281 and *cysT*– PA0282), and the C5-dicarboxylate transport (PA5530), all transporters are putative components of ABC transporters (PA2204, PA3019), and putative amino acid permeases (PA3641, PA0789). Additionally, three genes (a probable AGCS Na^+^/alanine/glycine symporter – PA2252, a probable amino acid permease – PA3641, and *nqrB* – PA2998) involved in Na^+^ ions transport (GO:0006814) were induced in the T48 variants (fold change 3.10, 2.42 and 2.11 respectively; Table S1 and S2).

There is also a set of genes (14 in total) coding for unknown functions that are differentially regulated, half of them are up-regulated and the rest down-regulated (Table 2). Interestingly, when these results are linked to the genome sequence analysis, none of the differentially regulated hypothetical proteins had mutations in the ORF or the promoter. One of these up-regulated hypothetical proteins is *yjjT* (PA4627), which product could be a putative rRNA (Guanin-N2)-methyltransferase (GO:0008990) according to the Gene Ontology analysis.

### Gene Ontology Analysis

A global analysis of the differentially regulated genes by Gene Ontology (GO) was performed. Such analysis revealed that the majority of the proteins are grouped according to the “catalytic activity” and “binding” biological functions (Figure 4A, Table S3). When analysed the cellular component category of the GOs, “cell part” which comprises the “plasma membrane” and “intracellular” categories and “macromolecular complex” represent the two groups (Figure 4B). Additionally, the regulated genes could be involved in two biological processes: “metabolic process” and “cellular process” (Figure 4C). When analysed, the product genes’ functions, “transferases”, “oxidoreductases”, “hydrolases”, and “lyases” are the most abundant protein functions (Figure 4D). Stressed bacteria followed a complex adaptive response that involves different biological processes such as the regulation of oxidation-reduction process, regulation of cell shape, transmembrane transport systems and cell redox homeostasis.

### Salt tolerance assay of *P. aeruginosa* mutants

The detection of a large number of genes differentially regulated in the T48 variant (Table 2) confirmed the hypothesis that high salt resistance in T48 is linked to many genes that participate together in the adaptive response of *P. aeruginosa*. However, it is difficult to determine the individual contribution of each gene in the adaptation to hypersalinity. For this reason, to investigate the individual contribution of each one of the differentially regulated genes in the T48 variant, we decided to explore their available mutants in the

*P. aeruginosa* PA14 transposon insertion library [21]. From 78 differentially regulated genes, mutants for 39 genes could be recovered from the library (Table S2). These individual knock-out mutants were tested for growth at different NaCl concentrations. Our results indicated that only mutants in *ccoO2* (cytochrome *c* oxidase subunit), PA4517 (conserved hypothetical protein), *dnaK* (chaperone), and *hslU* (ATP-dependent protease) showed a significant difference with the wild type when grown in 8.5 and 500 mM NaCl (Figure3). Some mutants (in PA1556, PA4517 and PA4761) grew worse than the parental strain in low salt concentration (8.5 mM NaCl), while all they did grow better in high salinity medium (500 mM).

## Discussion

The results of this study show that *P. aeruginosa* can survive and adapt to prolonged extreme stress conditions. The obtained data suggest that the differential response to salt stress between T0 and T48 variants is not linked to specific mutational events. This in line with previous finding with these clones recovered from salt, where has been shown that after several passages, bacteria recover their normal phenotype [14]. However, we cannot discard that some of the detected mutations could play a role in the observed phenotypes. Genetic manipulation of *P. aeruginosa* at single nucleotide level is nowadays still a challenge.

The gene expression analysis revealed that many genes are differentially regulated in the stressed cells. The differential induction of membrane transporters may reflect altered ion fluxes between the bacterial cell and the surrounding medium to maintain homoeostasis. In fact, the primary response of bacteria to a highly osmotic environment is the accumulation of certain solutes, like K^+^, glutamate, trehalose, proline, and glycine betaine, at concentrations that are proportional to the osmolarity of the medium [15].

When bacteria face a growing concentration of Na^+^, they actively transport K^+^ ions [15]. A recent study demonstrated that a steady K^+^ supply, even under unfavourable energetic conditions, plays a key role in long-term survival and desiccation tolerance for *Halobacteriumsalinarum* within salt crystal [16]. This is consistent with the fact that Na^+^/ K^+^ transporters were slightly induced in the T48 variant, including, the glutathione-regulated K^+^-efflux system protein KefB (PA1207; 1.5-fold regulation) and a putative K^+^ channel (PA1496; 1.52-fold regulation). K^+^ transporters are regulated by an increase in environment osmolarity regardless of the solute used and turgor. This response is modulated by the external concentrations of Na^+^. The K^+^ ions act as a cytoplasmic-signaling molecule, activating and/or inducing enzymes and transport systems that allow the cell to adapt to elevated salinity [12, 15].

Additionally, three genes involved in Na^+^ ion transport (GO:0006814) were induced in the T48 variants (PA2252, PA3641 *nqrB* – PA2998; fold change 3.10, 2.42 and 2.11 respectively; Tables S1 and S2). The product of *nqrB* gene is a unique energy-transducing complex, widely distributed among marine and pathogenic bacteria. It converts the energy from the oxidation of NADH and the reduction of quinone into an electrochemical Na^+^-gradient that can provide energy to the cell [17]. In addition, it allows the Na^+^ ion to pass through the hydrophobic core of the membrane and provides cation specificity to the translocation system [18]. These results are complementary with the 6 up-regulated and 13 down-regulated transporters (Table 2). From these 6 up-regulated transporters, *rnfC* (PA3491) is related with the electron transport complex, which is overexpressed when IscR is up-regulated [19]. In contrast, *rnfE* (PA3494), the putative periplasmic component of the RNF system [20], is underexpressed in T48 variants, a contradiction as the RNF system is very close to the Na^+^-pumping NADH:ubiquinone oxidoreductase [20, 21]. Another important up-regulated transporter is *czcB* (PA2521), which is associated with resistance to heavy metals [22–24], but recently it was discovered that it is also responsible for Ca^2+^ ions homeostasis [25]. Other up-regulated transporters are *narK1* (PA3877; related with the nitrate respiration under anaerobic conditions [25, 26]), *ompI* (PA3894; related to the aminoglycoside resistance [27]), and two more probable transporters with unknown associated metabolites (PA1876, *yhiH* – PA5231). Despite the fact that the gene coding for OmpI is up-regulated in T48 (Table 2), MexCD (PA4598 and PA4599) transporters, which are related to fluoroquinolones resistance [81, 82], are down-regulated. Similarly, LptG (PA3827), a lipopolysaccharide export system permease, is also down-regulated. Regarding transporters related to carbohydrate transport, RbsA (PA1947) and YhhS (PA1993) are also down-regulated. Finally, the rest of the down-regulated transporters are putative MFS and ABC type transporters of unknown metabolites, except for a putative K^+^ channel (PA1496) and YdfC (PA2777), a putative formic/nitrite transporter which was also found to be expressed under antibiotic stress [28, 29].

Four members of the 8-gene operon *iscR*-PA3808 the ferredoxin Fdx2 (PA3809), the co-chaperone HscB (PA3811), the L-cysteine desulfuraseIscS (PA3814), and the iron-sulphur cluster assembly transcription factor IscR (PA3815) are up-regulated, supporting previous observations about their expression in high concentration of salts [19]. Surprisingly, we did not find any significant up-regulation of the IscR-regulated ferredoxin FprB (PA4615), which are also associated with salt stress [30]. Other up-regulated genes are implicated in post-translational modifications such as *trmD* (PA3743; tRNA guanosine methyltransferase) and *endA* (PA2749; DNA-specific endonuclease I). Methylation of coding or non-coding RNA might play an important role in gene expression regulation [31]. Moreover, the S-adenosylmethionine decarboxylase proenzyme (*speD*; PA0654), involved in spermidine biosynthesis, was up-regulated in the T48 variants. Interestingly, previous works reported that spermidine is effective in alleviating the adverse effect of salt stress on plants [32, 33]. A recent finding indicated that spermidine priming treatments enhanced the antioxidant systems in plants exposed to salt stress and contributed to improved ion homeostasis [32–34]. Similarly, in *P. aeruginosa*, Johnson *et al* [34] reported that spermidine plays an important function as an organic polycation to bind lipopolysaccharide and to stabilize the cell surface. It protects the outer membrane from aminoglycoside antibiotics, antimicrobial peptides, and oxidative stress.

Another up-regulated gene coding for a two-component system response regulator PmrA (PA4776), was identified in this study, which was recently reported to be associated with polymyxin resistance and hence osmotic stress [35].

The *rsmA* gene, known to be a regulator of the secondary metabolism and a carbon storage regulator, is down-regulated in salt-tolerant clones (Table 2). RsmA was found to play a very important role in early pathogenesis, especially in early colonisation and dissemination [36], due to its relevance in the expression of Type VI Secretory System (T6SS) [37, 38]. Moreover, *rsmA*-knockouts strains of *P. aeruginosa* have altered the expression of genes involved in a wide variety of pathways, including iron acquisition, formation of multidrug efflux pumps and motility [39]. The genes coding for TssK1 (PA0079) and IcmF2 (PA1669) are also down-regulated. Both proteins are fundamental for the pathogenesis of *P. aeruginosa*, as the former is implied in the assembly of T6SS complex [40] while the latter is involved in the virulence [41].

Most of these T3SS proteins are members of the 12-gene operon *popN*-*popD*, which are expressed not only in pathogenesis but also under different environmental stresses such as low concentration of Ca^2+^ or direct contact with host cells [42]. Additionally, ExoS and ExoT have an ADP-ribosyltransferase activity, playing an important role in the bacterial survival and dissemination in clinical strains [43, 44, 45].

Salt stress also showed impact on three metabolic genes that were up-regulated: the cytochrome *c* oxidase (*coxA*; PA0106), a putative acyl-CoA dehydrogenase (PA0508), and the anthranilate-coenzyme A ligase (*pqsA*; PA0996). Previously, the putative acyl-CoA dehydrogenase, a gene associated with changes in membrane fluidity [46], was found overexpressed in *Burkholderia pseudomallei* when treated with NaCl. The fact that *pqsA* is up-regulated may indicate that the *Pseudomonas* quinolone signal (PQS) could be also overexpressed during salt encapsulation. This protein shapes bacterial population structure to survive under stressful environments and kills sensitive bacteria at a time that promotes anti-oxidative stress response [47].

When transcriptional regulators are analysed, only a putative transcriptional regulator (PA0547) is up-regulated in T48, having a potential role in the differential regulation of gene expression. However, *hslU* (PA5054) and *dnaK* (PA4761) genes, which encode for chaperone activity are found significantly down-regulated. This observation is in contrast with previous studies, where *dnaK* was overexpressed, being relevant in salt resistance in *Lactococcuslactis* [48]. Moreover, *dnaK* was also found up-regulated in marine bacteria allowing the adaptation to cold environments [49]. Possibly, these chaperones were up-regulated in salt-trapped bacteria but, once the T48 variant was recovered, these genes are quickly down-regulated due to other salt stress adaptations, the same maybe true for other genes.

CgrA (PA2127), which is found to be related to the expression of RsmN and, thus, the repression of RsmA [38], is up-regulated in T48 variant. The *cgrA* gene plays a key role in the expression of fimbrial genes and is related to MvaT mutants or anaerobic growth [50]. Despite the fact that only *narK1* (PA3877) was found to be up-regulated, results indicate that most of the up-regulated genes in T48 are associated to aerobic growth. Additionally, no mutations in the MvaT transcriptional regulator were revealed by whole genome analysis.

According to our results, salt resistance could be considered as a priming response, i.e. as a physiological process by which organisms prepare themselves for more quick or aggressive situations to future biotic or abiotic stress [51]. Although this phenomenon has been studied mostly in plants, there are also some examples of priming in the bacterial world. One critical issue is to explore how the signals that induce priming are received and transduced by the cells and prepare the bacteria for long-term persistence if growth is not possible. In plants, priming to salinity plays an important role as adaptive phenotypic strategy [52]. This process could develop different defence mechanisms in the cells against salinity stress such as antioxidant defence systems, the repair of membranes and the osmotic adjustment [53]. This kind of response is characterised, essentially, by the slow induction of many genes that together contribute to the acquisition of quick and effective adaptive strategy against stressor conditions. In such cases, molecular mechanisms responsible for priming effects are involved in the accumulation of signalling proteins or transcription factors [54], as well as epigenetic mechanisms [55, 56]. These epigenetic mechanisms are thought to bring a faster and more potent response to subsequent exposure to stress. This idea is supported for the Gene Ontology analysis, which suggested that some genes were involved in regulation of transcription, methylation process, response to stimulus, RNA metabolic processes and quorum sensing.

Interestingly, neither the mutants nor the wild-type were able to grow in DM with 1 M NaCl. All these genes showed a decreased transcription in the T48 variant and, consequently, if they are involved in adaptation to hypersalinity, a better growth under high NaCl concentration is expected when inactivated. Moreover, no sequence changes between the five genes from T0 and T48 were found, suggesting that the differences in growth under high salt conditions may be due to differential regulation, which requires further research to be clarified. The preservation of a long-lasting phenotype is not new in bacteria. For instance, the lactose metabolization response in *Escherichia coli* is maintained during more than ten generations after the removal of lactose due to the inheritance of very stable proteins [57].

## Conclusions

*Pseudomonas aeruginosa* can survive in inclusions of seawater crystals for many years. Upon recovery, this bacterium shows a better ability to grow in highly saline conditions, and the adaptation seems to be only phenotypic but not genetic, indicating a ‘priming’ phenomenon in this plastic bacterium. Although we have identified several genes potentially involved in adaptation to saline environments, the exact mechanisms which are responsible for priming in *P. aeruginosa* remain unclear. Our study provides a good start toward a deep understanding of the long-term salt stress behaviour of *P. aeruginosa*.

## Methods

### Bacterial model and growth conditions

*P. aeruginosa* ATCC 27853 (wild-type: T0 in this study) was grown overnight at 37°C in nutrient broth, centrifuged at 13,000 rpm for 10 min, washed three times and then suspended in filtered sterilise seawater to a final concentration of approximately 10^9^ CFU/ml in three independent replicas. Cells were incubated during fourteen years in closed Erlenmeyer flasks at room temperature. The concentration of salts in the used seawater was 37 g/l. The water was allowed to evaporate, the saline crystals were apparent after eight months, and the culture became completely desiccated after ten months. The initial number of CFU/ml was confirmed by serial decimal dilutions in nutrient agar.

#### Revitalisation of the bacterial cells

*P. aeruginosa* cells, maintained during fourteen years in sterilised seawater, were revitalised by the addition of 100 ml of sterilised nutrient broth to the salt crystal in the Erlenmeyer flask and incubated at 37°C with 100 rpm of shaking. Subsequent plating of an aliquot from this culture on nutrient agar yielded observable colonies. A few isolated colonies from the different replicas were recovered and saved for further analysis. Biochemical profiles of *P. aeruginosa* ATCC 27853 and the resuscitated cells (T48 variant in this study) were characterised using API 20NE system (bio-Merieux, France).

#### Evaluation of fitness in different salinity conditions

Bacterial growth curves were carried out in flat-bottomed 96-well microplates (Nunc, Denmark). Each well was filled with 100 μl of Davis Minimal medium (DM: Na2HPO4 6.78g/l; KH2PO4 3g/l; NaCl 0.5g/l; NH4Cl 1g/l; 1 mM MgSO4; 0.1 mM CaCl2, 0.28% Glucose and 0.25% casamino acids), supplemented with NaCl to final concentrations of 2M, 1M, 500mM, 250mM and 8.5mM. Overnight cultures of T0 and T48

*P. aeruginosa* cells were added to a final OD595 of 0.04. The growth of T0 and T48 variants was followed with four replicas of each one in the same concentrations of NaCl. Microplates were incubated in an Infinite F200 TECAN microplate reader for 24 hours at 37°C with 15s of shaking duration, 3mm of shaking amplitude. The interval time of absorbance measurements at 595nm was 15min. The same manipulation was repeated adding 100mM KCl to selected concentrations of NaCl.

#### Whole genome sequencing after recovery from salt

Libraries were prepared using a TruSeq DNA PCR-Free Library Preparation Kit (Illumina, USA) and were sequenced on anIllumina-MiSeq system using a 600-cycle v3 reagent kit, resulting in 300-bp paired-end reads. Sequence data are available from the NCBI database under Bioproject accession PRJNA420955. A reference genome for strain

*P. aeruginosa* ATCC 27853 was assembled using A5-miseq version 20140604 and annotated using prokka version 1.12-beta [59]. Snippy version 3.2 [60] was used to identify variants in strains c1-5 (clone1 to clone5) relative to the reference complete genome of *P. aeruginosa* ATCC 27853 (Genbank accession CP015117). Assembly of 1,818,724 error-corrected reads (estimated 47.68-fold coverage) resulted in 46 contigs with an N50 of 353 kb and a total size of 6.79 Mb.

#### Transcriptome profiling of salt-trapped *P. aeruginosa*

To find out what genes are involved in the differential salinity resistance, global transcription profile of cultures of *P. aeruginosa* T0 and its derivative T48 variant were carried out using microarray technology. Bacterial cells were grown overnight in Davis minimal medium (DM) at 37°C under 200 rpm of shaking. Three independent 1/50 dilutions of each of them were grown until they reached an optical density of 0.5 at 600 nm. The cells were washed and resuspended in DM supplemented with RNA protect reagent (Qiagen, Germany). Cell lysis and total RNA extractions were performed with the RNeasy mini kit according to the manufacturer’s recommendations (Qiagen, Chatsworth, CA), except that 1 mg/ml of lysozyme was used to lyse *Pseudomonas* cells. DNase digestions were carried out on the column by adding 82 units of Kunitz enzyme (Qiagen) with incubation at room temperature for 15 min. An additional DNase digestion was performed on the purified RNA to ensure the absence of DNA. RNA quality was checked through agarose electrophoresis before cDNA synthesis. Fluorescently labelled cDNA for microarray hybridisation was obtained by using the SuperScript Indirect cDNA Labelling System (Invitrogen) as recommended by the supplier. Briefly, 20 μg of total RNA was transformed to cDNA with Superscript III reverse transcriptase using random hexamers as primers and including aminoalyl-modified nucleotides in the reaction mixture. After cDNA purification, Cy3 or Cy5 fluorescent dye (Amersham Biosciences) was coupled to the amino-modified first-strand cDNA. The labelling efficiency was assessed by using a NanoDrop ND1000 spectrophotometer (NanoDrop Technologies). Equal amounts of Cy3- or Cy5-labelled cDNAs, one of them corresponding to the control and the other to the problem under analysis, were mixed and dried in a Speed-Vac. Labelled cDNA was hybridised to *P. aeruginosa* microarray slides version 2 from the Pathogen Functional Genomics Resource Center from J. Craig Venter Institute Microbial Hybridization of Labelled Probes protocol. Following hybridization, the slides were washed, dried, and scanned using a ScanArray Express scanner and software (Packard BioScienceBioChip Technologies). For the analysis of DNA microarray slides, background correction and normalization of expression data were performed using LIMMA [61]. To avoid the exaggerated variability of log ratios for low-intensity spots during local background correction, we used the *normexp* method in LIMMA to adjust the local median background estimates. The resulting log ratios were print-tip LOESS normalised for each array [62]. Only genes that exhibited changes compared to the wild-type control of two-fold and more, as well as P values of ≤0.05, were considered in the study. Finally, to explore the functional roles of the regulated genes, the Gene Ontology (GO) analysis was performed through the PANTHER online software [63] and QuickGO tool [64].

#### Salt tolerance assay of *P. aeruginosa* mutants

The desired mutants were isolated from PA14 transposon insertion mutants [65]. The selected *P. aeruginosa* PA14 mutants, with deletions in genes showing transcriptional variation in the microarray experiments, were used. Salt tolerance of these mutants was measured and compared to the wild-type strain PA14. The salt tolerance assay was performed on 96-well polystyrene plates. Each well was filled with 100 μl of DM minimal medium supplemented with NaCl to final concentrations of 8.5 mM (DM with no NaCl added), 250 mM and 500 mM, with four replicas for each NaCl concentration. The microplates were incubated at 37°C, and the optical density at 600 nm was measured after 24 hours. For mutants showing statistically significant differences in growth respect the wild-type strain on NaCl (>25%), salt tolerance assay was repeated for each sodium chloride concentration.

#### Statistical analysis

All parameters for the growth curves were estimated using Growthcurver [66]. Using this data, all model parameters —carrying capacity, initial population size, growth rate, doubling time and the empirical area under the curve—for all growth curves of both variants, T0 and T48, were compared using Student’s *t* test according to the different NaCl and KCl concentrations. Additionally, two-sided Kolmogorov-Smirnov tests were applied to compare the growth curves per treatments. *P* values less than or equal to 0.05 were considered statistically significant. All statistical tests were performed in R v. 3.4.4 [67].

## Acknowledgements

We are grateful to Prof. Jens Rolff for support and helpful comments from Yeliz Karatas, Arpita Nath from Freie Universität Berlin and Dr Paul Smith.

## Availability of data and materials

All data are available in the manuscript and supplementary material. The sequences and genomic data were deposited as indicated in the manuscript.

## Authors’ contributions

EH, EGT, CIQ, AB, KG and ARR carried out the experimental work. ARR and JB designed the experimental work. PJ contributed with genome sequencing and analysis. All authors conducted analytical work. EH and ARR drafted the manuscript with input from all authors. All authors read and approved the final manuscript.

## Ethics approval and consent to participate

Not applicable.

## Consent for publication

Not applicable.

---

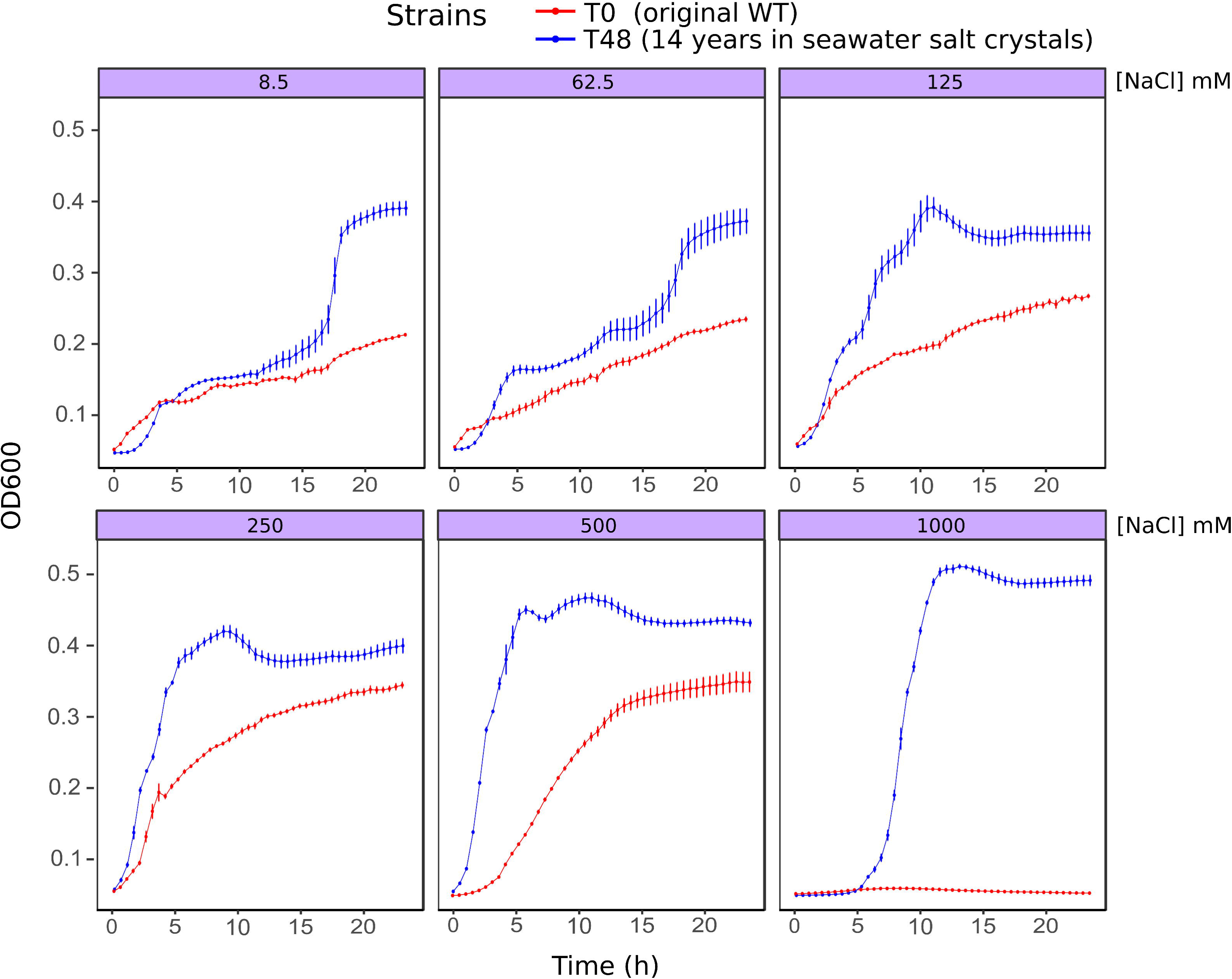

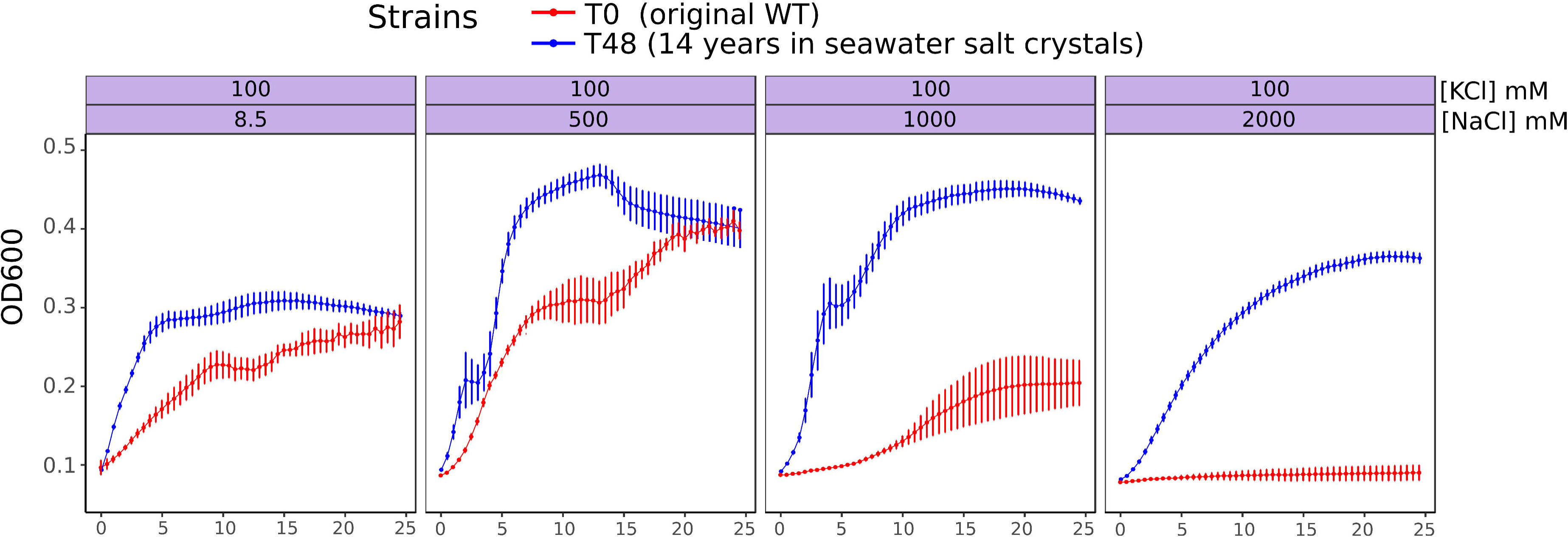

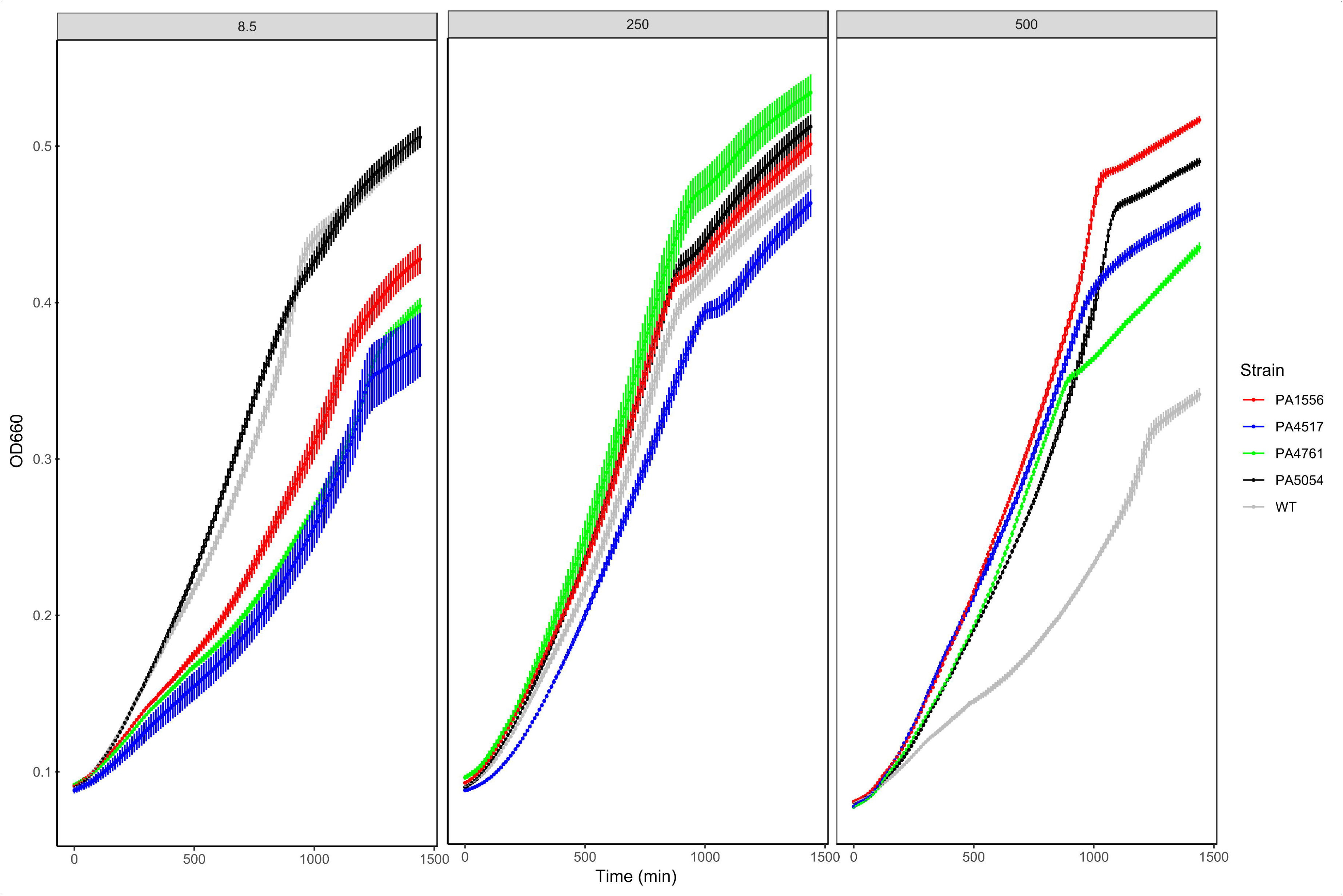

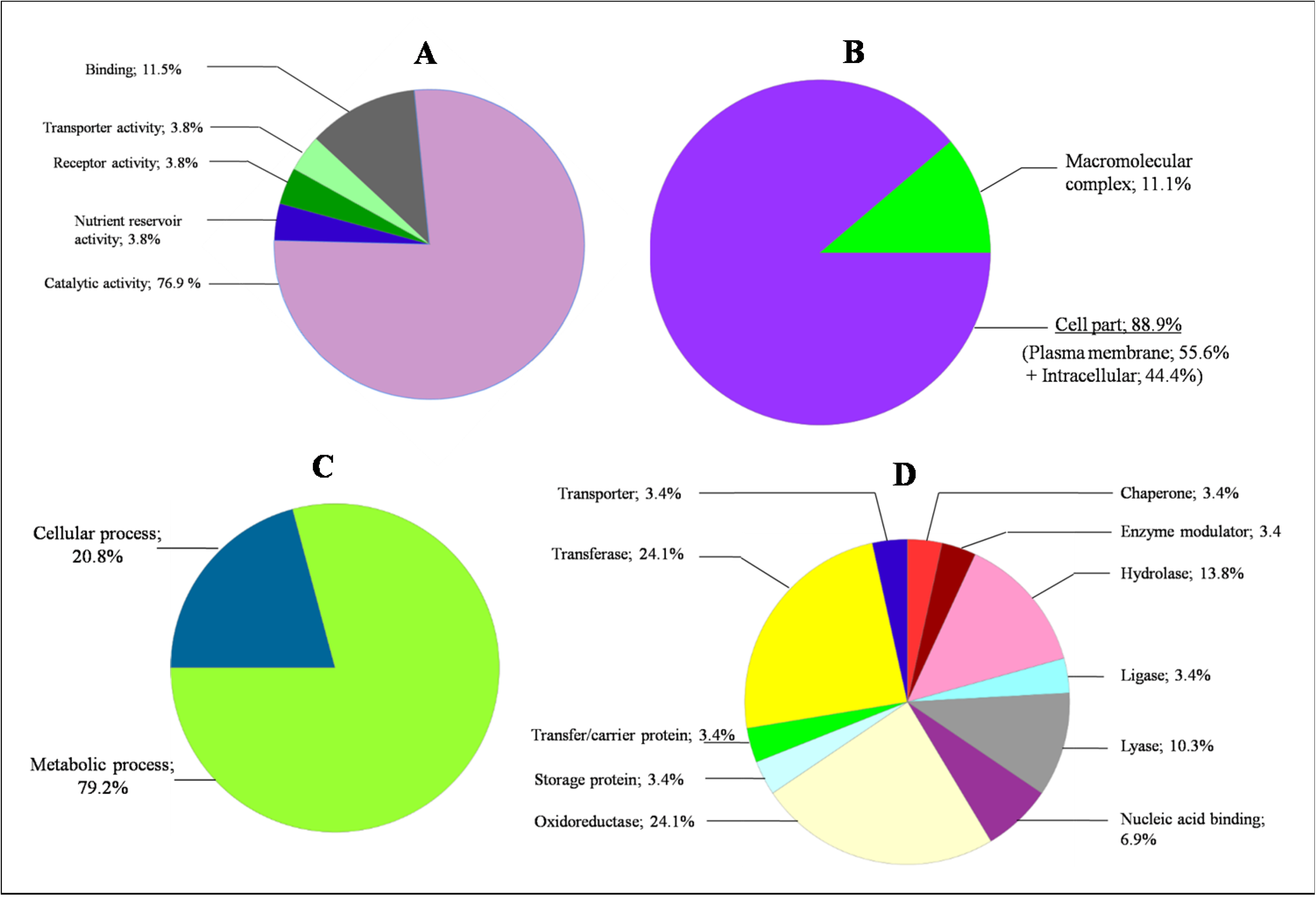

