## Supplementary material for "Seawater salt-trapped *Pseudomonas aeruginosa* survives for years and gets primed for salinity tolerance": table s1

Table S1. The detected mutations in *P. aeruginosa* clones ATCC27853 after Whole Genome Sequencing.

| **Reference Genome** | **Position** | **Reference nucleotide** | **Clone 1** | **Clone 2** | **Clone 3** | **Clone 4** | **Clone 5** | **WT** | **Locus tag** | **Gene product** | **Effect** |
|---|---|---|---|---|---|---|---|---|---|---|---|
| CP015117 | 1557965 | G | A | – | – | – | – | – | A4W92_07285 | aromatic amino acid transporter | missense_variant c.400C>T p.Leu134Phe |
| CP015117 | 2248912 | T | – | – | G | – | – | G | A4W92_10450 | prepilin-type N-terminal cleavage/methylation domain-containing protein | missense_variant c.211_212delTAinsGC p.Tyr71Ala |
| CP015117 | 2248913 | A | – | – | C | – | – | C | A4W92_10450 | prepilin-type N-terminal cleavage/methylation domain-containing protein | missense_variant c.211_212delTAinsGC p.Tyr71Ala |
| CP015117 | 3593639 | G | C | C | C | – | – | C | A4W92_16615 | DNA polymerase III subunit beta | synonymous_variant c.795G>C p.Arg265Arg |
| CP015117 | 3695540 | G | – | C | – | – | – | – | A4W92_17055 | FHA domain-containing protein | missense_variant c.338C>G p.Ala113Gly |
| CP015117 | 4059537 | C | T | T | – | – | – | T | A4W92_18780 | hybrid sensor histidine kinase/response regulator | missense_variant c.4204C>T p.Arg1402Cys |
| CP015117 | 4461847 | A | A | G | | | | G | intergenic | | |
| CP015117 | 5180188 | A | A | T | A | A | | A | intergenic | | |
| CP015117 | 5282094 | C | – | – | – | – | T | T | intergenic | | |
| CP015117 | 6810389 | T | – | C | C | – | – | C | A4W92_31605 | hypothetical protein | missense_variant c.16T>C p.Ser6Pro |
| CP015117 | 6810407 | T | – | C | C | – | – | C | A4W92_31605 | hypothetical protein | synonymous_variant c.34T>C p.Leu12Leu |
| CP015117 | 6810427 | C | – | – | T | – | – | T | A4W92_31605 | hypothetical protein | synonymous_variant c.54C>T p.Ser18Ser |
| CP015117 | 6810434 | T | – | T | G | – | – | G | A4W92_31605 | hypothetical protein | missense_variant c.61T>G p.Cys21Gly |

-Clone 1 to Clone 5: *P. aeruginosa* Biosamples available from NCBI database under accession numbers: SAMN08127309; SAMN08127310; SAMN08127311; SAMN08127312 and SAMN08127313.

-WT: wild-type strain
