## Supplementary material for "Seawater salt-trapped *Pseudomonas aeruginosa* survives for years and gets primed for salinity tolerance": table s2

Table S2. Gene list of mutants recovered from the *P. aeruginosa* PA14 mutant insertion library and used in this study to test the impact of salt in the growth.

| **Gene (Pseudomonas ID)** | **Description** |
|---|---|
| PA0310 | Hypothetical protein |
| PA0547 | Probable transcriptional regulator |
| *speD* (PA0654) | S-adenosylmethionine decarboxylase proenzyme |
| *rsmA* (PA0905) | Ribosomal RNA small subunit methyltransferase A |
| *yehS* (PA0915) | Conserved hypothetical protein |
| *shaD* (PA1057) | $Na^+/H^+$ antiporter subunit D |
| PA1209 | Hypothetical protein |
| *fixG* (PA1551) | Probable ferredoxin |
| *ccoP2* (PA1555) | cbb3-type cytochrome *c* oxidase, CcoP subunit |
| *ccoO2* (PA1556) | cbb3-type cytochrome *c* oxidase, CcoO subunit |
| *ccoN2* (PA1557) | cbb3-type cytochrome *c* oxidase, CcoN subunit |
| *pcr3* (PA1701) | YscX family type III secretion protein, preprotein translocase X |
| *pcrV* (PA1706) | Type III secretion protein PcrV |
| *pcrH* (PA1707) | Regulatory protein PcrH |
| *popB* (PA1708) | Translocator protein PopB |
| *popD* (PA1709) | Translocator outer membrane protein PopD precursor |
| *exsB* (PA1712) | Exoenzyme S synthesis protein B |
| PA2501 | Hypothetical protein |
| PA2662 | Conserved hypothetical protein |
| PA2757 | Hypothetical protein |
| *uup* (PA3019) | Probable ATP-binding component of ABC transporter |
| *gapA* (PA3195) | Glyceraldehyde 3-phosphate dehydrogenase |
| *bfrB* (PA3531) | Bacterioferritin |
| *trmD* (PA3743) | tRNA (guanine-N1)-methyltransferase |
| *iscS* (PA3814) | Pyridoxal phosphate-dependent L-cysteine desulfurase |
| *iscR* (PA3815) | Iron sulfur biosynthesis cluster operon transcriptional regulator IscR |
| *ispA* (PA4043) | Geranyltranstransferase |
| *oprG* (PA4067) | Outer membrane protein OprG precursor |
| PA4390 | Hypothetical protein |
| *cysD* (PA4443) | ATP sulfurylase small subunit |
| PA4517 | Conserved hypothetical protein |
| PA4611 | Hypothetical protein |

| | |
|---|---|
| *yjjT* (PA4627) | Conserved hypothetical protein |
| *ftsJ* (PA4752) | Cell division protein FtsJ |
| *dnaK* (PA4761) | Chaperone protein DnaK |
| *prmA* (PA4850) | Ribosomal protein L11 methyltransferase |
| *hslU* (PA5054) | Heat shock protein HslU |
| PA5174 | Probable beta-ketoacyl synthase |
| PA5530 | C5-dicarboxylate transporter |
