## Supplementary material for "Seawater salt-trapped *Pseudomonas aeruginosa* survives for years and gets primed for salinity tolerance": table s3

| **Gene ID** | **Gene Name<br>Gene Symbol** | **PANTHER<br>Family /Subfamily** | **PANTHER Protein Class** |
|---|---|---|---|
| **Up-regulated Genes** | | | |
| PA3641\|UniProtKB=<br>Q9HXZ0 | Probable amino acid permease<br>PA3641 | SUBFAMILY NOT NAMED<br>(PTHR30330:SF1) | - |
| PA0281\|UniProtKB=<br>Q9I6K9 | Sulfate transport protein CysW<br>*cysW* | Sulfate transport system permease protein CysW<br>(PTHR30406:SF1) | - |
| PA4850\|UniProtKB=<br>Q9HUW3 | Ribosomal protein L11 methyltransferase<br>*prmA* | Electron transfer flavoprotein beta subunit lysine methyltransferase<br>(PTHR43648:SF1) | DNA methyltransferase<br>RNA methyltransferase |
| PA5174\|UniProtKB=<br>Q9HU15 | Beta-ketoacyl-[acyl-carrier-protein] synthase FabY<br>*fabY* | 3-oxoacyl-[acyl-carrier-protein] synthase 1<br>(PTHR11712:SF306) | acyltransferase<br>dehydrogenase<br>esterase<br>ligase<br>methyltransferase |
| PA1708\|UniProtKB=<br>Q9I324 | Translocator protein PopB<br>*popB* | - | - |
| PA1709\|UniProtKB=<br>Q9I323 | Translocator outer membrane protein PopD<br>*popD* | - | - |
| PA1707\|UniProtKB=<br>Q9I325 | Regulatory protein PcrH<br>*pcrH* | - | - |
| PA1710\|UniProtKB=<br>P26995 | Exoenzyme S synthesis protein C<br>*exsC* | - | - |
| PA0547\|UniProtKB=<br>Q9I5Y9 | Probable transcriptional regulator<br>PA0547 | - | - |
| PA1228\|UniProtKB=<br>Q9I4B0 | Uncharacterized protein<br>PA1228 | - | - |
| PA0282\|UniProtKB=<br>Q9I6K8 | Sulfate transport protein CysT<br>*cysT* | Sulfate transport system permease protein CysT<br>(PTHR30406:SF8) | - |
| PA1714\|UniProtKB=<br>Q9I321 | ExsD<br>*exsD* | - | - |
| PA1701\|UniProtKB=<br>G3XD37 | Uncharacterized protein<br>PA1701 | - | - |
| PA2749\|UniProtKB=<br>Q9I094 | DNA-specific endonuclease I<br>*endA* | Endonuclease-1<br>(PTHR33607:SF2) | - |
| PA1719\|UniProtKB=<br>P95434 | Type III export protein PscF<br>*pscF* | - | - |
| PA4412\|UniProtKB=<br>Q9HW01 | UDP-N-acetylglucosamine--N-acetylmuramyl-(pentapeptide) pyrophosphoryl-undecaprenol N-acetylglucosamine transferase<br>*murG* | Glycosyltransferase<br>(PTHR21015:SF22) | acetyltransferase<br>glycosyltransferase<br>transfer/carrier protein |
| PA3814\|UniProtKB=<br>Q9HXI8 | Cysteine desulfurase IscS<br>*iscS* | Selenocysteine lyase<br>(PTHR11601:SF52) | lyase |
| PA1695\|UniProtKB= | Translocation protein in type III | - | - |

| | | | |
|---|---|---|---|
| Q9I332 | secretion<br>*pscP* | | |
| PA2252\|UniProtKB=<br>Q9I1L7 | Probable AGCS sodium/alanine/glycine symporter<br>PA2252 | SUBFAMILY NOT NAMED<br>(PTHR30330:SF1) | - |
| PA4043\|UniProtKB=<br>Q9HWY4 | Geranyltranstransferase<br>*ispA* | Geranylgeranyl pyrophosphate synthase 10, mitochondrial-related<br>(PTHR43281:SF1) | acyltransferase |
| PA5429\|UniProtKB=<br>Q9HTD7 | Aspartate ammonia-lyase<br>*aspA* | Aspartate ammonia-lyase<br>(PTHR42696:SF2) | lyase |
| PA1057\|UniProtKB=<br>Q9I4R7 | Uncharacterized protein<br>PA1057 | SUBFAMILY NOT NAMED<br>(PTHR34584:SF1) | - |
| PA1319\|UniProtKB=<br>Q9I425 | Cytochrome bo(3) ubiquinol oxidase subunit 3<br>*cyoC* | Cytochrome bo(3) ubiquinol oxidase subunit 3<br>(PTHR11403:SF2) | oxidase |
| PA3182\|UniProtKB=<br>Q9X2N2 | 6-phosphogluconolactonase<br>*pgl* | 6-phosphogluconolactonase<br>(PTHR11054:SF0) | hydrolase |
| PA3815\|UniProtKB=<br>Q9HXI7 | IscR<br>*iscR* | HTH-type transcriptional regulator IscR<br>(PTHR33221:SF10) | - |
| PA3841\|UniProtKB=<br>G3XDA1 | Exoenzyme S<br>*exoS* | SUBFAMILY NOT NAMED<br>(PTHR10339:SF30) | - |
| PA2757\|UniProtKB=<br>Q9I086 | Uncharacterized protein<br>PA2757 | - | - |
| PA3019\|UniProtKB=<br>Q9HZI7 | Probable ATP-binding component of ABC transporter<br>PA3019 | ABC transporter ATP-binding protein uup<br>(PTHR19211:SF69) | - |
| PA2991\|UniProtKB=<br>P57112 | Soluble pyridine nucleotide transhydrogenase<br>*sthA* | Soluble pyridine nucleotide transhydrogenase<br>(PTHR22912:SF93) | dehydrogenase<br>oxidase<br>reductase |
| PA1838\|UniProtKB=<br>Q9I2Q7 | Sulfite reductase<br>*cysI* | Sulfite reductase [NADPH] subunit beta<br>(PTHR11493:SF47) | - |
| PA3809\|UniProtKB=<br>Q51383 | 2Fe-2S ferredoxin<br>*fdx* | 2Fe-2S ferredoxin<br>(PTHR23426:SF34) | - |
| PA3131\|UniProtKB=<br>Q9HZ93 | Probable aldolase<br>PA3131 | KHG/KDPG aldolase<br>(PTHR30246:SF0) | aldolase |
| PA2662\|UniProtKB=<br>Q9I0H6 | Uncharacterized protein<br>PA2662 | - | - |
| PA1699\|UniProtKB=<br>G3XCT8 | Uncharacterized protein<br>PA1699 | - | - |
| PA1700\|UniProtKB=<br>Q9I329 | Uncharacterized protein<br>PA1700 | - | - |
| PA2998\|UniProtKB=<br>Q9HZK7 | Na(+)-translocating NADH-quinone reductase subunit B<br>*nqrB* | SUBFAMILY NOT NAMED<br>(PTHR30578:SF1) | - |
| PA0044\|UniProtKB=<br>Q9I788 | Exoenzyme T<br>*exoT* | SUBFAMILY NOT NAMED | - |

| | | | |
|---|---|---|---|
| | | (PTHR10339:SF30) | |
| PA4442\|UniProtKB=O50274 | Bifunctional enzyme CysN/CysC<br>*cysNC* | Elongation factor 1-alpha 1-related<br>(PTHR23115:SF170) | G-protein<br>hydrolase<br>translation elongation factor<br>translation initiation factor |
| PA5012\|UniProtKB=G3XD35 | Heptosyltransferase II<br>*waaF* | ADP-heptose--LPS heptosyltransferase 2<br>(PTHR30160:SF7) | carbohydrate kinase<br>glycosyltransferase |
| PA2204\|UniProtKB=Q9I1R3 | Probable binding protein component of ABC transporter<br>PA2204 | - | - |
| PA1318\|UniProtKB=Q9I426 | Cytochrome bo(3) ubiquinol oxidase subunit 1<br>*cyoB* | Cytochrome bo(3) ubiquinol oxidase subunit 1<br>(PTHR10422:SF35) | oxidase |
| PA3811\|UniProtKB=Q9HXJ1 | Co-chaperone protein HscB homolog<br>*hscB* | Co-chaperone protein HscB<br>(PTHR14021:SF16) | - |
| PA4002\|UniProtKB=G3XD88 | Rod shape-determining protein<br>*rodA* | Rod shape-determining protein RodA<br>(PTHR30474:SF1) | - |
| PA0915\|UniProtKB=Q9I542 | Uncharacterized protein<br>PA0915 | SUBFAMILY NOT NAMED<br>(PTHR37805:SF1) | - |
| PA0284\|UniProtKB=Q9I6K6 | Uncharacterized protein<br>PA0284 | - | - |
| PA1706\|UniProtKB=G3XD49 | Type III secretion protein PcrV<br>*pcrV* | - | - |
| PA1712\|UniProtKB=P26994 | Exoenzyme S synthesis protein B<br>*exsB* | - | - |
| PA4627\|UniProtKB=Q9HVG4 | Ribosomal RNA small subunit methyltransferase C<br>*rsmC* | Ribosomal RNA small subunit methyltransferase C<br>(PTHR18895:SF70) | DNA methyltransferase<br>RNA methyltransferase |
| PA0789\|UniProtKB=Q9I5E9 | Probable amino acid permease<br>PA0789 | SUBFAMILY NOT NAMED<br>(PTHR43341:SF2) | - |
| PA1717\|UniProtKB=Q9I318 | Type III export protein PscD<br>*pscD* | - | - |
| PA5530\|UniProtKB=Q9HT43 | Probable MFS dicarboxylate transporter<br>PA5530 | SUBFAMILY NOT NAMED<br>(PTHR43528:SF5) | - |
| PA1696\|UniProtKB=Q9I331 | Translocation protein in type III secretion<br>*pscO* | - | - |
| PA3743\|UniProtKB=Q9HXQ1 | tRNA (guanine-N(1)-)-methyltransferase<br>*trmD* | tRNA (guanine-N(1)-)-methyltransferase<br>(PTHR32125:SF1) | - |
| PA1723\|UniProtKB=Q9I314 | Type III export protein PscJ<br>*pscJ* | SUBFAMILY NOT NAMED<br>(PTHR30046:SF2) | - |

| | | | |
|---|---|---|---|
| PA3195\|UniProtKB=P27726 | Glyceraldehyde-3-phosphate dehydrogenase<br>*gap* | glyceraldehyde-3-phosphate dehydrogenase C-related (PTHR43148:SF2) | dehydrogenase |
| PA4390\|UniProtKB=Q9HW14 | Uncharacterized protein<br>PA4390 | - | - |
| PA1697\|UniProtKB=Q9I330 | ATP synthase in type III secretion system<br>PA1697 | SUBFAMILY NOT NAMED (PTHR15184:SF48) | ATP synthase<br>DNA binding protein<br>anion channel<br>hydrolase<br>ligand-gated ion channel |
| PA0654\|UniProtKB=Q9I5R7 | S-adenosylmethionine decarboxylase proenzyme<br>*speD* | S-adenosylmethionine decarboxylase proenzyme (PTHR33866:SF1) | - |
| PA4443\|UniProtKB=O50273 | Sulfate adenylyltransferase subunit 2<br>*cysD* | Sulfate adenylyltransferase subunit 2 (PTHR43196:SF1) | nucleotidyltransferase |
| PA3068\|UniProtKB=Q9HZE0 | NAD-specific glutamate dehydrogenase<br>*gdhB* | SUBFAMILY NOT NAMED (PTHR43403:SF1) | dehydrogenase |
| PA0570\|UniProtKB=Q9I5W6 | Uncharacterized protein<br>PA0570 | - | - |
| **Down-regulated genes** | | | |
| PA1557\|UniProtKB=Q9I3G0 | Cytochrome c oxidase, cbb3-type, CcoN subunit<br>*ccoN2* | SUBFAMILY NOT NAMED (PTHR10422:SF29) | oxidase |
| PA1555\|UniProtKB=Q9I3G2 | Cbb3-type cytochrome c oxidase subunit<br>*ccoP2* | SUBFAMILY NOT NAMED (PTHR33751:SF1) | - |
| PA3531\|UniProtKB=Q9HY79 | Ferroxidase<br>*bfrB* | Bacterioferritin (PTHR30295:SF0) | storage protein |
| PA5446\|UniProtKB=Q9HTC1 | Uncharacterized protein<br>PA5446 | - | - |
| PA4752\|UniProtKB=P95454 | Ribosomal RNA large subunit methyltransferase E<br>*rlmE* | rRNA methyltransferase 2, mitochondrial (PTHR10920:SF18) | - |
| PA0310\|UniProtKB=Q9I6I1 | Uncharacterized protein<br>PA0310 | SUBFAMILY NOT NAMED (PTHR12907:SF22) | - |
| PA3572\|UniProtKB=Q9HY48 | Uncharacterized protein<br>PA3572 | - | - |
| PA4611\|UniProtKB=Q9HVH9 | Uncharacterized protein<br>PA4611 | - | - |
| PA0905\|UniProtKB=O69078 | Carbon storage regulator homolog<br>*csrA* | Carbon storage regulator (PTHR34984:SF1) | - |
| PA4067\|UniProtKB=Q9HWW1 | Outer membrane protein OprG<br>*oprG* | Outer membrane protein W (PTHR36920:SF1) | - |
| PA1556\|UniProtKB=Q9I3G1 | Cytochrome c oxidase, cbb3-type, CcoO subunit<br>*ccoO2*<br>ortholog | - | - |
| PA5054\|UniProtKB= | ATP-dependent protease ATPase | ATP-dependent protease | chaperone |

| | | | |
|---|---|---|---|
| Q9HUC5 | subunit HslU<br>hslU<br>ortholog | ATPase subunit HslU<br>(PTHR43815:SF1) | |
| PA1551\|UniProtKB=<br>Q9I3G6 | Probable ferredoxin<br>PA1551<br>ortholog | SUBFAMILY NOT<br>NAMED<br>(PTHR24960:SF45) | - |
| PA2501\|UniProtKB=<br>Q9I0Y1 | Uncharacterized protein<br>PA2501<br>ortholog | - | - |
| PA1209\|UniProtKB=<br>Q9I4C9 | Uncharacterized protein<br>PA1209<br>ortholog | SUBFAMILY NOT<br>NAMED<br>(PTHR42709:SF7) | - |
| PA4761\|UniProtKB=<br>Q9HV43 | Chaperone protein DnaK<br>dnaK<br>ortholog | Stress-70 protein,<br>mitochondrial<br>(PTHR19375:SF184) | - |
| PA4517\|UniProtKB=<br>Q9HVQ5 | Uncharacterized protein<br>PA4517<br>ortholog | Phosphoethanolamine<br>transferase EptC<br>(PTHR30443:SF2) | - |
