## Supplementary figures and images for "Seawater salt-trapped *Pseudomonas aeruginosa* survives for years and gets primed for salinity tolerance"

### figure s1

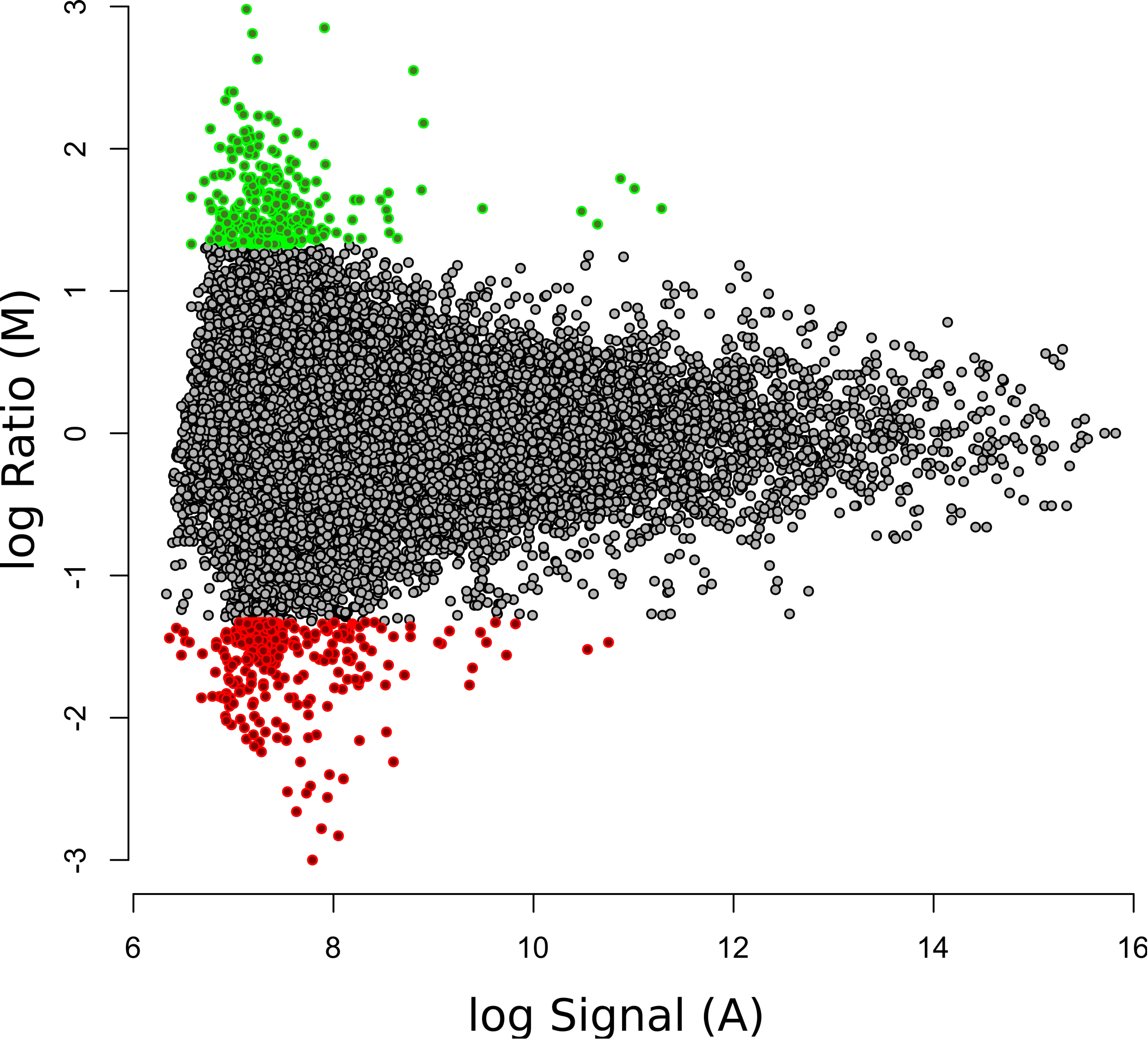
